# Oxytocin and Vasopressin Immunoreactivity Differs Across Auditory Brainstem Nuclei in Rodents with Distinct Social Systems

**DOI:** 10.64898/2026.09.28.755085

**Authors:** Luberson Joseph, Lucas Boren, Grace Ingram, Elizabeth A. McCullagh

## Abstract

Oxytocin (OT) and vasopressin (AVP) are neuropeptide hormones involved in regulating animal social behavior and a broad spectrum of physiological processes. Although their distributions are well documented in neuroendocrine regions of the forebrain and midbrain, their expression in the hindbrain remains poorly understood. Here, we used immunohistochemistry to quantify OT and AVP immunoreactive puncta within three auditory brainstem nuclei, the lateral superior olive (LSO), the medial superior olive (MSO), and the medial nucleus of the trapezoid body (MNTB) in six wild-caught rodent species differing in sociality. We also quantified the volume of these nuclei and examined variation in total brain volume across species and sociality. OT and AVP puncta count differed among species and social groups. Group-living species exhibited higher OT and AVP puncta counts than monogamous and solitary species in the LSO and MNTB. In the MSO, OT puncta counts did not differ among social groups, whereas AVP puncta counts were higher in group-living than in monogamous and solitary species. Total brain volume and the volumes of the MNTB and MSO differed among species, but not across social groups, whereas LSO volume did not differ among species or sociality. These findings revealed sociality-related variation in OT and AVP immunoreactive puncta within auditory brainstem circuits and suggest that neuropeptide signaling within early auditory brainstem pathways may contribute to the neural integration of social and auditory information.

## INTRODUCTION

Comparative analyses provide a powerful framework in biological research for dissecting the diversity of social behavior and the neural pathways that support them. This approach moves beyond single-species characterization and enables the identification of conserved and divergent mechanisms that shape social organizations and life-history strategies across both vertebrates and invertebrates. For example, mammalian species that evolved solitary lifestyles generally display a high degree of aggression toward conspecifics, rely exclusively on maternal care, and maintain non-overlapping home ranges (Makuya & Schradin, 2024b, 2024a; Šklíba et al., 2009). In contrast, group-living species tend to exhibit greater social affiliation and tolerance, provide maternal care, share overall home-ranges, and display selective aggression toward unfamiliar individuals (Ebensperger & Cofré, 2001; Krause & Ruxton, 2002). Monogamous or pair-living species further differ by forming enduring pair bonds, engaging in biparental care, and jointly defending territories (Lukas & Clutton-Brock, 2013; Wittenberger & Tilson, 1980). These structural differences in social behavior strategies are thought to be the product of evolutionary pressures arising from ecological constraints, resources distribution, mate competition, and predation risks (Xu et al., 2010). These may also be accompanied by species-specific differences in central nervous system organization and perhaps in variation in neuromodulator hormones associated with social behavior in the brain.

Rodents comprise nearly 40% of all described mammalian species and represent an exceptional model system for comparative studies of social behavior (Joseph, et al., 2026; Solari & Baker, 2007). Their extensive diversity in social strategies, ranging from solitary to highly social, and eusocial-living, make them particularly well suited for investigating the neural bases of species-typical social strategies. Extensive comparative research has sought to identify the neural mechanisms underlying variation in social behavior across rodent taxa (Bester-Meredith & Marler, 2001; Xu et al., 2010). These studies provide strong evidence that interspecific differences in social behavior are associated with differences in the density and distribution of central neuromodulators across distinct brain regions. A classic example is the work by Bester-Meredith and Marler (2001), who showed that the highly aggressive and promiscuous white-footed mice (*Peromyscus leucopus*) exhibited higher vasopressin (AVP) fiber densities in basal forebrain regions such as the bed nucleus of the stria terminalis, the amygdala, and the supraoptic nucleus compared to the less aggressive, monogamous California mouse (*Peromyscus californicus*). In addition, social Brandt’s voles (*Lasiopodomys braindtii*) exhibited greater densities of oxytocin (OT) and AVP immunoreactive cells in the lateral hypothalamus, the medial preotic area and the medial amygdala compared to the solitary greater long-tailed hamster (*Tscherskia triton*) (Xu et al., 2010). Similarly, studies in voles have demonstrated that experimental manipulation of OT, dopamine, and AVP neuromodulators can directly influence pair bonding, selective affiliation and other social behaviors in closely related species with different mating strategies (Aragona et al., 2006; Lim and Young, 2006). Together, these findings demonstrate that evolutionary variation in social behavior is accompanied by corresponding differences in neuromodulatory circuitry, suggesting that changes in the distribution and activity of neuromodulatory systems may contribute to the diversification of species-typical social phenotypes.

OT and AVP are nonapeptide hormones synthesized in magnocellular neurons of the supraoptic and paraventricular nuclei of the hypothalamus. Both neuropeptides play central roles in regulating social behaviors (Campbell, 2010; Froemke & Young, 2021) and numerous physiological processes (Hermesch et al., 2024; Uvnäs-Moberg et al., 2019; Wakerley et al., 1978). OT has been extensively characterized for its roles in parturition and lactation (Hashimoto et al., 1985). Beyond its peripheral effects, OT also modulates a broad spectrum of social behaviors, including social bond formation (Johnson & Young, 2015; Rigney et al., 2022), social motivation (Gordon et al., 2011), group cohesion (De Dreu et al., 2010; Samuni et al., 2017), social memory (Rimmele et al., 2009; Zhan et al., 2024), social avoidance (Duque-Wilckens et al., 2020), peer partner preferences (Goodwin et al., 2025), social novelty preferences (Smith et al., 2017; Tops et al., 2013), and maternal behavior (Mota-Rojas et al., 2023; Witchey et al., 2024). In contrast, AVP is well known for its functions in maintaining water homeostasis and arteriole contractions (Hashimoto et al., 1985), modulating male courtship behavior (Lim and Young, 2006), affiliative interactions (Carter, 1998), and aggression (Everts et al., 1997; Ferris et al., 1987). Given their broad and context-dependent roles, many studies have examined OT and AVP’s cellular and receptor expression across the forebrain and midbrain regions of species with different social systems (Xu et al. 2010; Johnson and Young, 2015; Rigney et al., 2022).

Despite decades of research on these neuropeptides’ role in regulating social behavior, direct cross-species comparisons of OT and AVP immunoreactive distribution in the hindbrain, specifically, in the lateral superior olivary complex (LSO), the medial superior olivary complex (MSO), and the medial nucleus of the trapezoid body (MNTB) remain scarce (Barberis & Tribollet, 1996; Kanwal & Rao, 2002). This gap is particularly notable in the auditory brainstem, where nuclei such as the LSO, MSO, and MNTB are essential for sound localization and auditory processing (Haragopal & Winters, 2023; Joseph, et al., 2025; Joseph, New, et al., 2025; Middlebrooks, 2015; Tollin, 2003) and to mediate social interaction, mate finding, and predator avoidance. These functions are integral in social communication, and we previously showed that group-living and monogamous rodents exhibit greater auditory sensitivity and have a more sensitive neural response to binaural stimuli than solitary living rodents (McCullagh et al. 2025). Accordingly, integrating comparative neuroanatomy to quantify the density of OT and AVP in the LSO, MSO, and MNTB of rodents that exhibit different social behavior strategies could be a step toward understanding how sensory processing circuits may be modulated to support species-specific social behavior and communication strategies.

In the present study, we used immunohistochemical and microscopy techniques to compare OT and AVP immunoreactive puncta in three main regions of the auditory brainstem (LSO, MSO, and MNTB) in six wild-caught rodent species that differ in social behavior strategies (solitary, monogamous, and group-living). We also measured brain anatomical features (brain length, brain width, and brain height) across social systems to explore the relationship between brain volume and social behavior. Because OT and AVP are critical in regulating social behavior, we hypothesized that rodents with more complex social behavior strategies (group-living and monogamous) will exhibit more immunoreactive puncta for OT and AVP in their LSO, MSO, and MNTB compared to solitary-living rodents. We also predicted that social rodents would exhibit larger brain volume and heavier brain mass compared to solitary rodents, due to increased cognitive demand associated with social interactions and group-living.

## MATERIALS AND METHODS

### Animals

Experiments were conducted on 30 male individuals representing six wild-caught rodent species (Table 1, N = 5 individuals per species), with males selected to maintain a consistent sex composition across species and to minimize potential sex-related variation in OT and AVP neuroanatomy. These animals represented a subset of individuals previously captured for an auditory brainstem responses (ABR) study (McCullagh et al., 2025). In brief, rodents were captured in seven locations in Oklahoma and Kansas between June 2022 and July 2025 using aluminum Sherman non-folding traps (3 in. x 3 in. x 10 in.) (H.B Sherman Traps, Inc., Tallahassee, FL) baited with oat and peanut butter (See McCullagh et al., 2025 for detailed trapping techniques and sampling locations). Animals were euthanized with a lethal dose of pentobarbital (120 mg/kg), perfused transcardially with phosphate buffered saline (PBS), followed by 4% paraformaldehyde (PFA) or 10% formalin. Animal brains were extracted from the skull and post-fixed overnight in fixative at 4°C. The brains were next transferred to PBS the following day and stored at 4°C until further immunohistochemical processing. All animal collection, handling, and procedures were conducted in accordance with regulations of the United States, the Oklahoma Department of Wildlife Conservation (ODWC), the Kansas Department of Wildlife and Parks (KDWP), the American Society of Mammalogists (Sikes et al., 2019), and the Oklahoma State University Institutional Animal Care and Use Committee (IACUC) in Stillwater (protocol number: 22-09).

**Table 1:** Rodent species included in the study and their social behavior classification. Common and scientific species name, family, and social behavior systems of all six species used in this study. Rodents were captured from six locations in Oklahoma including James Collin wildlife management area (WMA), Packsaddle WMA, Sandy Sanders WMA, Stillwater, Tulsa, and Selman living laboratory and one location from Kansas (Kansas University field station).

| Common name | Scientific name | Family | Social structure | References |
| --- | --- | --- | --- | --- |
| House mouse | *Mus musculus* | Muridae | Group-living | (Noyes et al., 1982) |
| Norway rat | *Rattus norvegicus* | Muridae | Group-living | (Schweinfurth, 2020) |
| Prairie voles | *Microtus ochrogaster* | Cricetidae | Monogamous | (Getz et al., 1993) |
| Northern grasshopper mouse | *Onychomys leucogaster* | Cricetidae | Monogamous | (Ruffer, 1965) |
| Hispid pocket mouse | *Chaetodipus hispidus* | Heteromyidae | Solitary-living | (Paulson, 1988) |
| Ord’s kangaroo rat | *Dipodomys ordii* | Heteromyidae | Solitary-living | (Langford, 1983) |

### Immunohistochemistry staining

Brains were sectioned coronally at 100 micrometers (μm) using a Leica VT1000S vibratome (Leica Biosystems Nussloch GmbH Heidelberger Strasse, Nussloch, Germany). For each animal, 9 to 16 free floating sections containing the LSO, MSO, and MNTB were selected and atlas-matched to the mouse brain atlas (Paxinos & Franklin, 2019). Selected sections were immediately placed in a blocking solution containing AB media (0.2 M phosphate buffer, 5 M sodium chloride, 10% triton, Millipore water, Bovine serum albumin) and 10% normal goat serum (NGS) and incubated on a shaker at room temperature for 1 hour. Primary and secondary antibodies were diluted in AB media and 10% NGS. Sections were incubated in primary antibodies overnight, covered with parafilm and aluminum foil on a shaker at 4°C, followed by 3 x 10-minute washes in PBS. Sections were next incubated in secondary antibodies for 1 hour at room temperature on a shaker. Sections were subsequently incubated in 1:100 fluorescent Nissl solution (Invitrogen™ NeuroTrace™ 435/455 Blue Fluorescent Nissl Stain, catalog # N21479, Thermo Fisher Scientific, Eugene, OR, USA) for 30 minutes. After staining, sections were transferred to phosphate buffer (PB) and mounted onto premium superfrost^TM^ microscope slides (Thermo Fisher scientific, Eugene, OR, USA), and cover-slipped with microscope cover glass with Fluoromount-G™ Mounting Medium (Cat. No. 00-4958-02, Thermo Fisher Scientific, Carlsbad, CA, USA).

### Antibody characterization

The rabbit polyclonal OT antiserum and guinea pig polyclonal AVP antiserum (see Table 2 for antibodies details) were used at a 1:500 concentration to detect OT and AVP immunoreactivity, respectively. The specificity of the OT antiserum has been demonstrated by antigen reabsorption, which eliminated staining in rat hypothalamic tissue following preincubation with OT (Constantinescu et al., 2026). The AVP antiserum was generated against a synthetic peptide corresponding to amino acids 24-32 of mouse vasopressin neurophysin 2-copeptin and has been validated for immunohistochemical detection of AVP in mouse and rat tissues (Fuyuki et al., 2026; Gu et al., 2025; Kuske et al., 2024). Consistent with previous applications (Constantinescu et al., 2026; Fuyuki et al., 2026; Gu et al., 2025; Kuske et al., 2024), the antibodies produce labeling similar to previous literature for OT and AVP immunoreactivity in the rodent LSO, MSO, and MNTB. Secondary antibodies (goat anti rabbit IgG, Alexa Fluor 647 and goat anti guinea pig IgG (H + L), Alexa Fluor 555, see Table 2 for antibodies details) were used at a 1:500 concentration to conjugate the primary antibodies with a fluorophore. Both secondary types are common in similar applications. Control experiments with secondary only application at the same concentration yielded no immunofluorescence suggesting specificity of signal to the primary antibodies.

**Table 2:** Antibodies used for immunohistochemistry. Primary and secondary antibodies used for immunohistochemical detection of oxytocin and vasopressin in the LSO, MSO, and MNTB of the auditory brainstem. Clonality, host, RRID, catalog number, company, and location.

| Antibody name | Type | Clonality | Host | RRID | Catalog # | Company | Location |
| --- | --- | --- | --- | --- | --- | --- | --- |
| Oxytocin antiserum | Primary | Polyclonal | Rabbit | AB_572258 | 200,68 | Immunostar | Wisconsin, USA |
| Vasopressin antiserum | Primary | Polyclonal | Guinea pig | AB_2725758 | 403,004 | SySy | Goettingen, Germany |
| Anti-Rabbit IgG (H+L),<br>Alexa Fluor™ 647 | Secondary | Polyclonal | Goat | AB_2535812 | A21244 | ThermoFisher<br>Scientific | Oregon, USA |
| Anti-Guinea Pig IgG<br>(H+L), Alexa Fluor™ 555 | Secondary | Polyclonal | Goat | AB_2535856 | A21435 | ThermoFisher<br>Scientific | Oregon, USA |

### Imaging and immunoreactive puncta count

Brain sections were imaged at 20x magnification and 0.41 pixel / μm using a ZEISS LSM 980 inverted Airyscan2 confocal microscope (Zeiss, Jena, Germany) under the 405, 555, and 647 lasers. Nissl bodies helped with anatomical identification of the three main nuclei of the auditory brainstem, including the LSO, MSO, and MNTB, based on the mouse brain atlas (Paxinos & Franklin, 2019). All images used to quantify OT and AVP immunoreactive puncta were analyzed in ImageJ/FIJI (Schindelin et al., 2012). For each image, the background was reduced by using rolling ball subtraction in ImageJ/FIJI. Next, the Huang method was used (AVP = 0 – 55, OT = 0 – 35) to convert images to grayscale and for thresholding. The watershed algorithm was used to separate individual immunoreactive puncta in each measured nuclei, and particles were analyzed by setting size from 2 to infinite (μm^2^) and circularity from 0.3 to 1 to manually quantify AVP and OT puncta (Figure 1). Quantification of OT and AVP puncta was performed on the entire field of view of the image taken for each brainstem region (LSO, MSO, and MNTB). For species (*Rattus norvegicus* and *Dipodomys ordii*) in which the MSO and MNTB were sometimes too large to be captured in a single 20x image, two separate images were acquired, and the measurements from those images were combined for the corresponding region.

**Figure 1:**
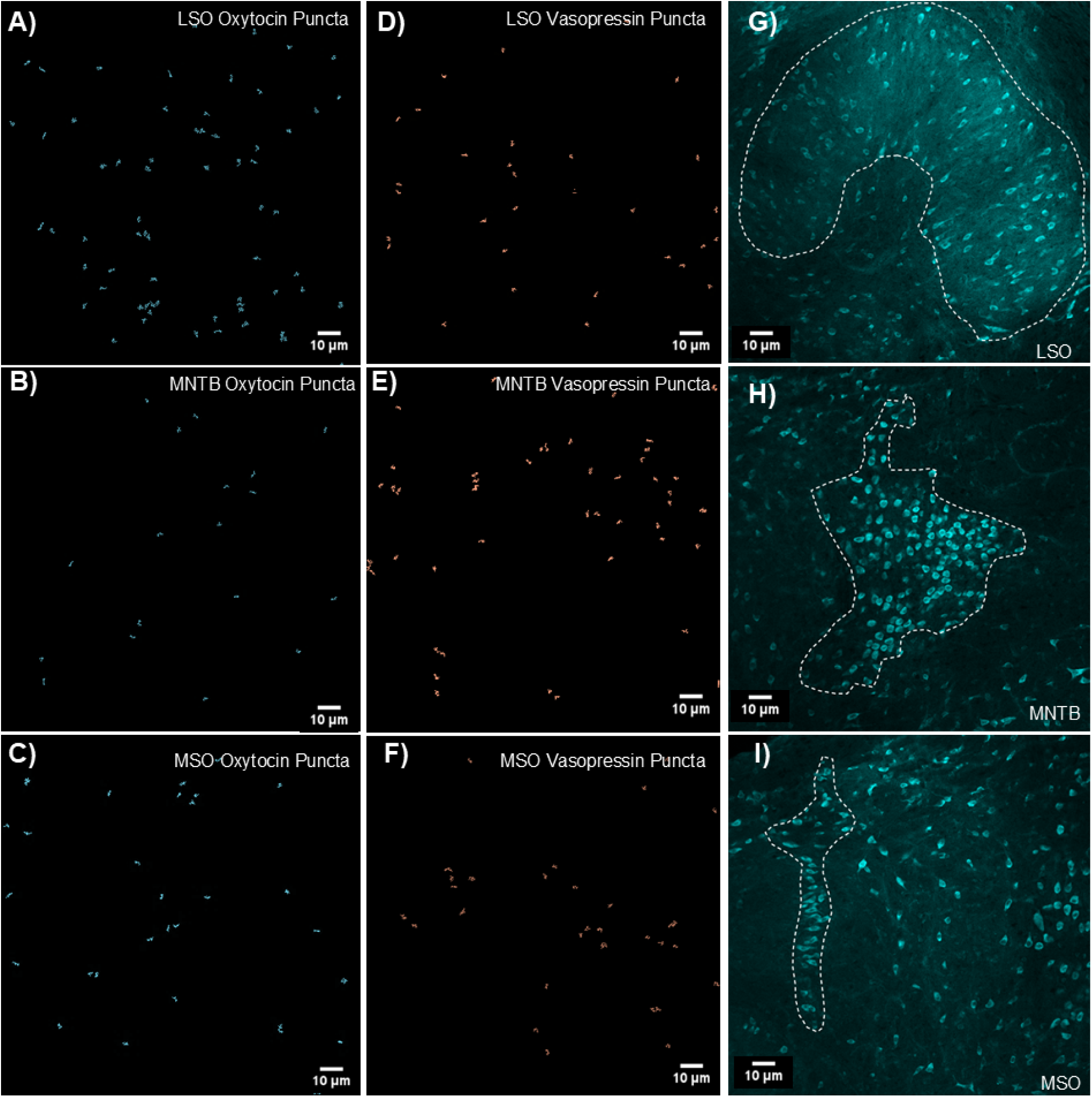
Oxytocin and AVP immunoreactive puncta and identification of auditory brainstem nuclei. A-C, representative images showing OT immunoreactive puncta in the LSO (A), MNTB (B), and MSO (C) of the prairie voles (*Microtus ochrogaster*). D-F, representative images showing AVP immunoreactive puncta in the LSO (D), MNTB (E), and MSO (F) of *M. ochrogaster*. G-I, representative Nissl-stained sections showing the anatomical location and boundaries of the LSO (G), MNTB (H), and MSO (I) in *M. ochrogaster*. Each immunoreactive puncta image was acquired at 20x magnification, while each of the anatomical location images were acquired at 10x magnification. White outlines indicate the boundaries of each nucleus used for volume measurements. See Supplementary Figure 1 for representative anatomical locations of the MNTB, MSO, and LSO of all species included in this study.

A six-inch stainless steel electronic vernier caliper (DIGI-Science Accumatic Digital Caliper Gyros Precision Tools, Monsey, New York, United Stated) was used to measure total brain length (L), brain width (W), and brain height (H) of all tested individuals to obtain a rough estimate of total brain volume of each species. Total brain length was defined as the distance from the tip of the olfactory bulbs to the tip of the medulla oblongata. Brain width was measured at the widest lateral point of the cerebral hemispheres. To determine brain height, the brain was positioned laterally and dorsoventral height was recorded at the tallest points of the cerebral hemisphere. These three measurements were used to calculate total brain volume of each animal using the ellipsoid formula: *V=(LxWxH)π/6* (Ma et al., 2025). Body mass and brain mass were measured for each animal using a digital stainless steel electronic scale (Weighmax W-2809 90 LB X 0.1 OZ, Durable Stainless Steel Digital Postal Scale, Chino, CA, USA).

Volumes of nuclei (MNTB, MSO, LSO) were measured in a separate cohort of individuals from those used in immunohistochemistry staining. However, all animals were collected from the same sampling sites during the same periods to ensure comparability. In total, 12 brainstems (two per species) were Nissl stained as described above and imaged at 10x using a ZEISS LSM 980 inverted Airyscan2 confocal microscope. The entire extent of the three nuclei (MNTB, MSO, and LSO) were imaged and manually outlined in Fiji/ImageJ as previously described (E. A. McCullagh et al., 2022). The cross-sectional area of each nucleus was measured using the measure function in FIJI/ImageJ. The volume of each nucleus was estimated by multiplying the area measured in each section by the section thickness (100 µm) and summed to get the total volume for the nucleus (Table 2). Nucleus volume was acquired for the left and right side of each nucleus for each animal, resulting in six measurements per section (two MNTBs, two MSOs, and two LSOs). All analyses were conducted on unadjusted images only.

**Table 2:** Descriptive statistics of volume measurement (mm^3^) for the six species (N is number of animals, n = number of sections per nucleus)

| Species | Social System | Nuclei volume in $\text{mm}^3$ (mean ± SE) | N (n) |
| --- | --- | --- | --- |
| *Chaetodipus hispidus* | Solitary | MNTB ($0.282 \pm 0.07$) | 2 (13-14) |
| | | MSO ($0.152 \pm 0.03$) | 2 (13-13) |
| | | LSO ($0.608 \pm 0.14$) | 2 (14-16) |
| *Dipodomys ordii* | Solitary | MNTB ($0.706 \pm 0.07$) | 2 (14-16) |
| | | MSO ($0.318 \pm 0.03$) | 2 (12-13) |
| | | LSO ($0.998 \pm 0.14$) | 2 (12-15) |
| *Microtus ochrogaster* | Monogamous | MNTB ($0.305 \pm 0.07$) | 2 (10-12) |
| | | MSO ($0.108 \pm 0.03$) | 2 (10-11) |
| | | LSO ($0.656 \pm 0.14$) | 2 (14-15) |
| *Onychomys leucogaster* | Monogamous | MNTB ($0.395 \pm 0.07$) | 2 (15-15) |
| | | MSO ($0.135 \pm 0.03$) | 2 (9-10) |
| | | LSO ($1.013 \pm 0.14$) | 2 (18-20) |
| *Mus musculus* | Group-living | MNTB ($0.258 \pm 0.07$) | 2 (13-14) |
| | | MSO ($0.031 \pm 0.03$) | 2 (6-7) |
| | | LSO ($0.749 \pm 0.14$) | 2 (14-15) |
| *Rattus norvegicus* | Group-living | MNTB ($0.503 \pm 0.07$) | 2 (13-15) |
| | | MSO ($0.162 \pm 0.03$) | 2 (12-12) |
| | | LSO ($1.004 \pm 0.14$) | 2 (15-16) |

### Statistical Analyses

All statistical analyses were conducted in R version 4.0.3 (R Core Team, 2020) with figures generated using the “ggplot2” package (Wickham, 2016). Histological measurements, including immunoreactive puncta counts in the MNTB, LSO, and MSO were analyzed using linear mixed-effects models (LMMs). For these analyses, nuclei puncta counts were modeled as the response variable, with species or sociality included as a fixed effect, and individual ID included as a random effect to account for repeated measurements within animal. Separate LMMs were fitted to evaluate the effects of species or sociality groups on puncta counts for each nucleus.

Next, linear models (LMs) were used to explore differences among species and sociality groups in brain volume and brain mass. Log-transformed brain volume or log-transformed brain mass was modeled as the response variable, species or sociality groups was included as a fixed effect. For both the LMM and LM models, body mass was log-transformed, mean-centered, and included as a continuous covariate to account for differences in individual body sizes within the datasets. When significant species or sociality effects were detected, estimated marginal means were calculated using the emmeans packages (Lenth, 2023), and used for pairwise comparisons among species or sociality groups. Multiple comparisons were adjusted using Tukey’s Honestly Significant Difference (HSD) method. For all models, residual diagnostic plots (Q-Q plots) were used to evaluate model assumptions and indicated no major deviations from normality.

In addition, the volume of the LSO, MSO, and MNTB, as well as the MNTB/LSO, MNTB/MSO, and MSO/LSO ratios, were analyzed using one-way analysis of variance (ANOVA), followed by Tukey’s HSD post hoc pairwise comparisons when significant effects were detected. Data are displayed as mean ± standard error (SE). Statistical significance was denoted as * p < 0.05, ** p ≤ 0.01, *** p ≤ 0.001.

## RESULTS

### Oxytocin immunoreactive puncta differ across species in the MNTB and LSO, but not in the MSO

We compared the number of OT immunoreactive puncta in the MNTB, LSO, and MSO in six wild-caught rodent species spanning diverse social structures, with log-transformed body mass included as a covariate to account for interspecific differences in body size. LMM revealed significant species differences in OT puncta counts in the MNTB (LMM: F_5, 30.20_ = 3.82, p = 0.008), and LSO (LMM: F_5, 30.41_ = 5.88, p < 0.001, supplementary figure 2). However, Tukey-adjusted pairwise comparisons did not identify significant differences among any of the six individual species in either the MNTB or the LSO. Log-transformed body mass was not significantly associated with OT-immunoreactive puncta counts in either the MNTB (F_1, 30.02_ = 3.11, p = 0.087) or the LSO (F_1, 30.85_ = 3.36, p = 0.076). In the MSO, neither species (F_5, 29.01_ = 1.92, p = 0.121) nor log transformed body mass (F_1, 27.12_ = 1.72, p = 0.200) was significantly associated with OT immunoreactive puncta counts. Accordingly, post hoc pairwise comparisons were not conducted for the MSO.

### Oxytocin immunoreactive puncta counts differ across social groups in the MNTB and LSO, but not in the MSO

Oxytocin immunoreactive puncta count also significantly differed across social groups in the MNTB (LMM: F_2, 30.54_ = 11.34, p < 0.001). Post-hoc comparisons indicated that group-living rodent species had significantly higher numbers of OT puncta counts than both monogamous (t-value = -3.826, p = 0.001) and solitary-living species (t-value = -3.956, p = 0.001, Figure 2A). However, OT puncta counts did not differ between monogamous and solitary species (t-value = 0.064 p = 0.997). Log-transformed body mass was significantly associated with MNTB OT immunoreactive puncta counts across sociality groups (LMM: F_1, 30.64_ = 6.31, p = 0.017). Similarly, OT immunoreactive puncta counts also significantly differ across sociality groups in the LSO (LMM: F_2, 30.24_ = 11.57, p < 0.001). Group-living species exhibited significantly higher OT puncta than both monogamous (t-value = -3.700, p = 0.002) and solitary-living species (t-value = -4.122, p < 0.001, Figure 2B). There was no difference in LSO OT puncta between monogamous and solitary-living species (t-value = -0.250, p = 0.966). Log-transformed body mass was also significantly associated with LSO OT immunoreactive puncta counts across social groups (LMM: F_1, 30.66_ = 5.68, p = 0.023). No significant differences in OT immunoreactive puncta counts were detected in the MSO across social groups (LMM: F_2, 30.72_ = 2.92, p = 0.068, Figure 2C). Because there were no social group effects in OT puncta counts in the MSO, pairwise comparisons were not performed among social groups. Log-transformed body mass was not significantly associated with MSO OT puncta counts across social groups (LMM: F_1, 31.84_ = 2.24, p = 0.144).

**Figure 2:**
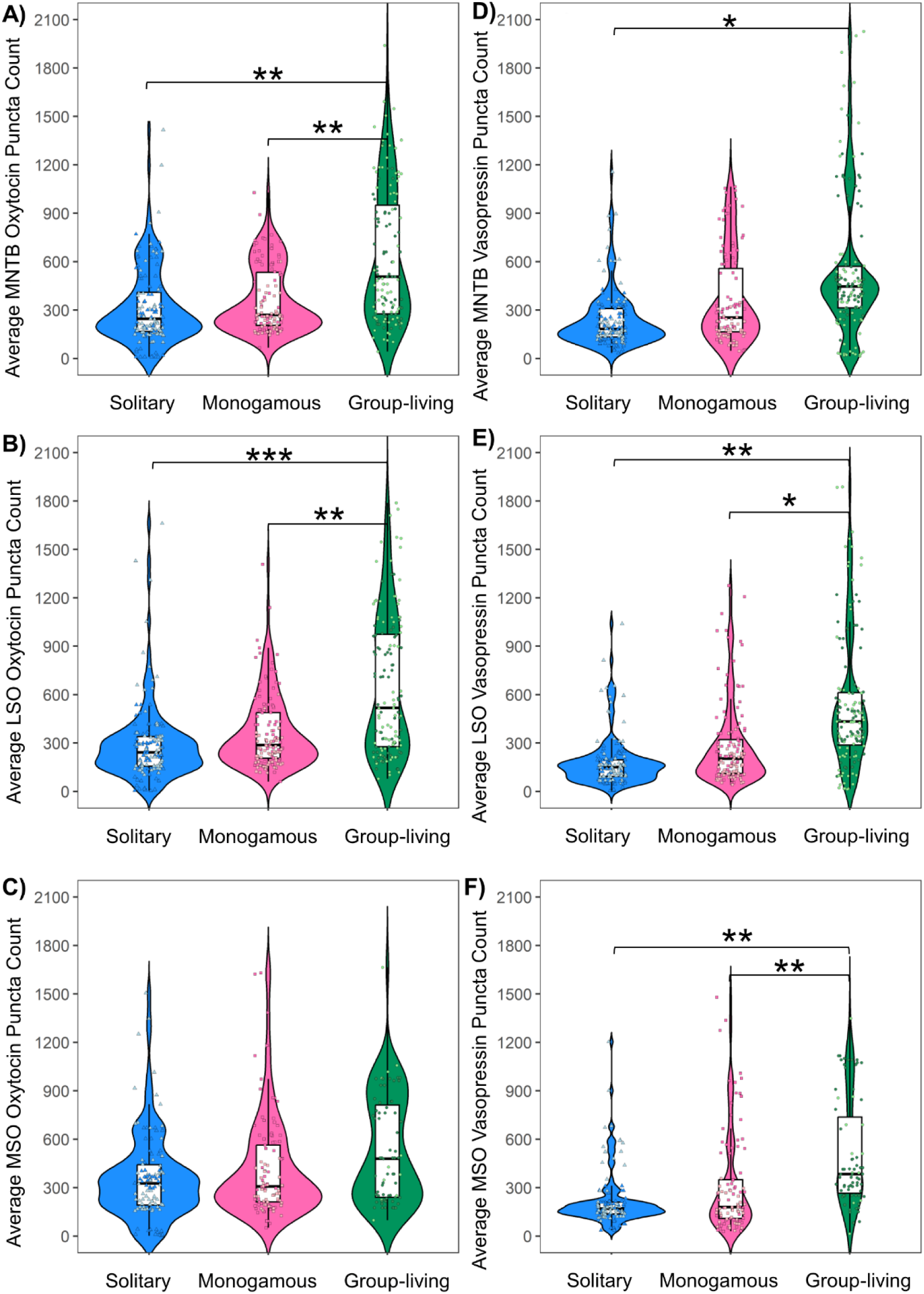
Oxytocin and AVP immunoreactive puncta count across social groups in the MNTB, LSO, and MSO. A-C, Average OT immunoreactive puncta counts across solitary (dodger blue), monogamous (hot pink), and group-living species (green) in the MNTB (A), LSO (B) and MSO (C), respectively. D-F, Average AVP-immunoreactive puncta counts across social groups in the MNTB (D), LSO (E), and MSO (F), respectively. Violin plots illustrate the distribution and density of puncta counts within each social group, while the embedded boxplots show the median and interquartile range; whiskers extend to 1.5 times the interquartile range. Individual point represent puncta count measurements and color according to species (dodger blue = *D. ordii*, light blue = *C. hispidus*, hot pink = *O. leucogaster*, light pink = *M. ochrogaster*, green = *R. norvegicus*, and light green = *M. musculus*), allowing visualization of variation both within and among species belonging to each social group. Significant differences in OT and AVP puncta counts were detected among species and social groups. Asterisks denoted statistically significant with * p < 0.05, ** p ≤ 0.01, *** p ≤ 0.001.

### Vasopressin immunoreactive puncta counts differ across species in the MNTB, LSO, and the LSO

LMM revealed significant species effects in AVP immunoreactive puncta counts in the MNTB (LMM: F_5, 30.19_ = 3.21, p = 0.019, supplementary figure 2). However, Tukey adjusted pairwise comparisons did not identify significant differences among individual species (all p > 0.05). Log-transformed body mass was not significantly associated with AVP immunoreactive puncta counts in the MNTB (LMM: F_1, 30.23_ = 0.76, p = 0.387). Similarly, AVP immunoreactive puncta counts differed significantly among species in the LSO (F_5, 30.29_ = 4.87, p = 0.002) and MSO (LMM: F_5, 29.78_ = 3.48, p = 0.013). However, Tukey adjusted pairwise comparisons did not identify significant differences among any individual species in either the LSO or the MSO (all p > 0.05). Log-transformed body mass was not significantly associated with AVP immunoreactive puncta counts in the LSO (LMM: F_1, 30.63_ = 2.95, p = 0.095) and the MSO (LMM: F_1, 28.55_ = 2.94, p = 0.096).

### Vasopressin immunoreactive puncta counts differ across social groups in the MNTB, LSO, and MSO

Vasopressin immunoreactive puncta counts significantly differed among social groups in the MNTB (F_2, 30.07_ = 4.46, p = 0.019). Post-hoc comparisons showed that group-living species exhibited significantly higher AVP immunoreactive counts than solitary species (t-value = - 2.716, p = 0.027, Figure 2D). However, no significant differences were detected either between monogamous species and solitary species (t-value = -0.656, p = 0.790), or between group-living species and monogamous species (t-value = -1.977, p = 0.133). Log-transformed body mass was not significantly associated with AVP puncta across social groups in the MNTB (F_1, 30.12_ = 0.25, p = 0.617). AVP-immunoreactive puncta also significantly differed among social groups in the LSO (F_2, 30.14_ = 8.17, p = 0.001). Tukey adjusted pairwise comparisons revealed that group-living species had significantly greater AVP puncta counts than monogamous (t-value = -2.780, p = 0.022) and solitary species (t-value = -3.633, p = 0.002, Figure 2E). AVP puncta counts did not differ between monogamous and solitary species (t-value = -0.724, p = 0.751) in the LSO. Log-transformed body mass was not significantly associated with AVP puncta across social groups in the LSO (F_1, 30.45_ = 1.99, p = 0.168). In addition, AVP immunoreactive puncta counts also differed across social groups in the MSO (F_2, 30.80_ = 5.63, p = 0.008). Group-living species exhibited higher AVP immunoreactive puncta counts than both solitary (t-value = -3.027, p = 0.012) and monogamous species in the MSO (t = -2.363, p = 0.019, Figure 2F). No significant differences were detected in MSO AVP puncta counts between monogamous and solitary species (t-value = -0.564, p = 0.840). Log-transformed body mass was not significantly associated with AVP puncta counts across social groups in the MSO region of the brainstem (LMM: F_1, 31.39_ = 2.18, p = 0.149).

### Total brain mass differs across species and social groups

Linear models showed significant differences in log-transformed brain mass among species (LM: Df = 5, F = 185.53, p < 0.001). Pairwise comparisons indicated that *D. ordii* had heavier brain mass than *C. hispidus* (p < 0.001), *M. ochrogaster* (p < 0.001), *M. musculus* (p < 0.001), and *O. leucogaster* (p < 0.001, supplementary figure 3). *R. norvegicus* also exhibited heavier brain mass than *C. hispidus* (p < 0.001), *M. ochrogaster* (p < 0.001), *M. musculus* (p < 0.001), and *O. leucogaster* (p < 0.001). *C. hispidus* (p < 0.001), *O. leucogaster* (p < 0.001), and *M. ochrogaster* (p = 0.005) had heavier brain mass than *M. musculus*. No significant differences were detected among the remaining species (all p > 0.05, supplementary table 1). Log-transformed body mass was not significantly associated with brain mass across species (Df = 1, F = 3.21, p = 0.086). Similarly, we detected a significant effect of sociality on log-transformed brain mass (LM: Df = 2, F = 11.31, p = 0.002, Figure 3A). Solitary species exhibited significantly heavier brain mass than monogamous species (t-value = -2.976, p = 0.016). However, no significant brain mass differences were observed either between group-living and monogamous (t-value = 0.565, p = 0.839) or between group-living and solitary species (t-value = -2.450, p = 0.054). Log-transformed body mass was significantly associated with brain mass across sociality groups (Df = 1, F = 96.004, p < 0.001).

**Figure 3:**
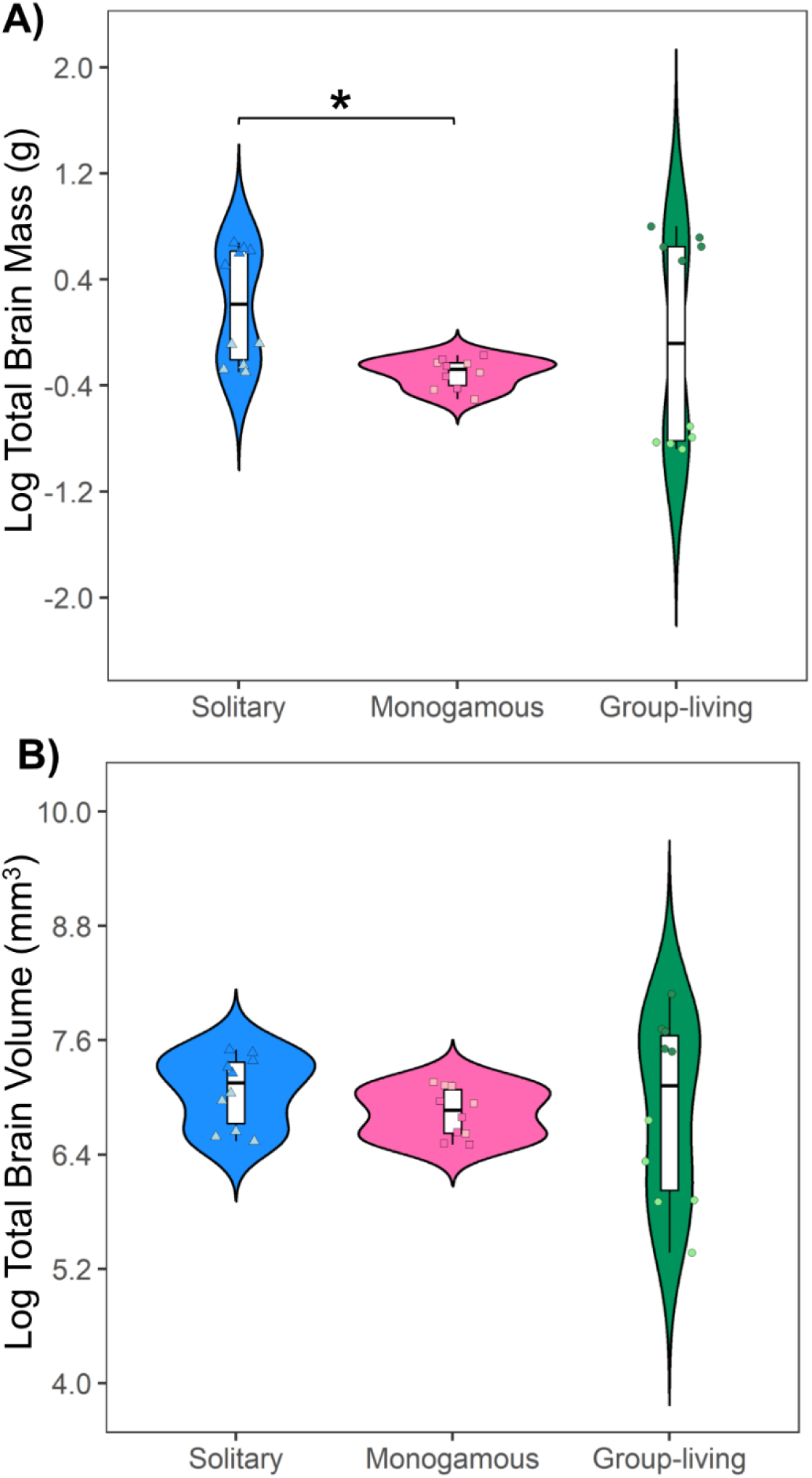
Log-transformed total brain mass and total brain volume across sociality groups. A) Log-transformed brain mass differences across solitary (dodger blue), monogamous (hot pink), and group-living species (green) species. B) Log-transformed brain volume across sociality groups. Violin plots illustrate the distribution and density of rodents’ log-transformed brain mass or brain volume within each sociality group, while the embedded boxplots show the median and interquartile range; whiskers extend to 1.5 times the interquartile range. Individual point represent brain mass and brain volume measurements and color according to species (dodger blue = *D. ordii*, light blue = *C. hispidus*, hot pink = *O. leucogaster*, light pink = *M. ochrogaster*, green = *R. norvegicus*, and light green = *M. musculus*), allowing visualization of variation both within and among species belonging to each sociality group. Significant differences log-transformed brain mass between solitary and monogamous rodents. No significant differences in log-transformed brain volume across sociality groups. Asterisks denoted statistically significant with * p < 0.05.

### Total brain volume differs across species, but not across social groups

We detected significant main effect of species on total brain volume (LM: Df = 5, F = 19.93, p < 0.001, supplementary figure 3). Tukey-adjusted pairwise comparisons showed that *M. musculus* exhibited significantly smaller brain volume than *D. ordii* (t-value = 4.101, p = 0.005), *M. ochrogaster* (t-value = 3.766, p = 0.011), and *R. norvegicus* (t-value = -3.654, p = 0.014). No significant pairwise differences were identified in brain volume among the remaining species (all p > 0.05, supplementary table 2). Log-transformed body mass was not significantly associated with log-transformed brain volume across species (LM: DF = 1, F = 0.255, p = 0.617). When comparing across social groups, the linear model revealed no significant effect of sociality on brain volume (LM: Df = 2, F = 1.237, p = 0.306, Figure 3B). Because there were no main effects of sociality on log-transformed total brain volume, pairwise comparisons were not performed. Log-transformed body mass was significantly associated with total brain volume across sociality groups (Df = 1, F = 52.05, p < 0.001). To account for interspecific variation in overall brain sizes, brain volume, and brain to body mass ratios were calculated for each individual. Neither log-transformed brain volume nor brain to body mass ratio differed significantly among sociality groups (supplementary figure 4: brain volume: Df = 2, F = 0.11, p = 0.897; brain mass: Df = 2, F = 2.42, p = 0.108). Because neither measure showed a significant overall effect of sociality, Tukey-adjusted pairwise comparisons were not performed.

### Brainstem nuclei volumes differ across species, but not across sociality groups

ANOVA revealed significant differences in MNTB (F = 5.078, p = 0.036) and MSO (F = 5.914, p = 0.025) volume across species (supplementary figure 5). Pairwise comparisons indicated that *D. ordii* had significantly larger MNTB (t-ratio = 4.163, p = 0.041) and MSO (t-value = 5.232, p = 0.014) than *M. musculus*. No significant differences in MNTB or MSO volume were detected among the remaining species (supplementary Table 3 and 4, respectively). In contrast, LSO volume did not differ across species (F = 1.719, p = 0.264). Because the overall ANOVA was not significant for LSO volume, post hoc pairwise comparisons were not performed among species. When species were grouped by sociality, no significant differences were detected in the volume of the MNTB (F = 0.649, p = 0.545), MSO (F = 3.126, p = 0.093), or LSO (F = 0.083, p = 0.921; Figure 4A). As no evident differences were detected for LSO, MNTB or MSO volumes across sociality groups, post hoc pairwise comparisons were not performed. To account for interspecific variation in overall nucleus size and to evaluate the relative proportions of the brainstem nuclei, MNTB/LSO, MNTB/MSO, and MSO/LSO ratios were calculated as previously described (McCullagh et al., 2022). A ratio of 1 indicates equal nucleus sizes, whereas values greater than 1 indicate that the numerator nucleus is larger than the denominator nucleus, and value less than 1 indicates that the numerator nucleus is smaller. No significant species differences were detected in either MNTB/LSO (F = 2.07, p = 0.202), MNTB/MSO (F = 2.144 p = 0.19) or MSO/LSO ratio (F = 1.84, p = 0.239, supplementary figure 6). However, when species were grouped by sociality, the MSO/LSO ratio differed significantly among social groups (F = 4.783, p = 0.038, Figure 4B). Post hoc comparisons indicated significant differences in the MSO/LSO ratio between solitary and group-living species (p = 0.038), reflecting a proportionally larger LSO relative to the MSO in solitary species. In contrast, neither the MNTB/LSO (F = 3.005, p = 0.100) nor the MNTB/MSO ratios (F = 2.051, p = 0.184) differed significantly among social groups.

**Figure 4:**
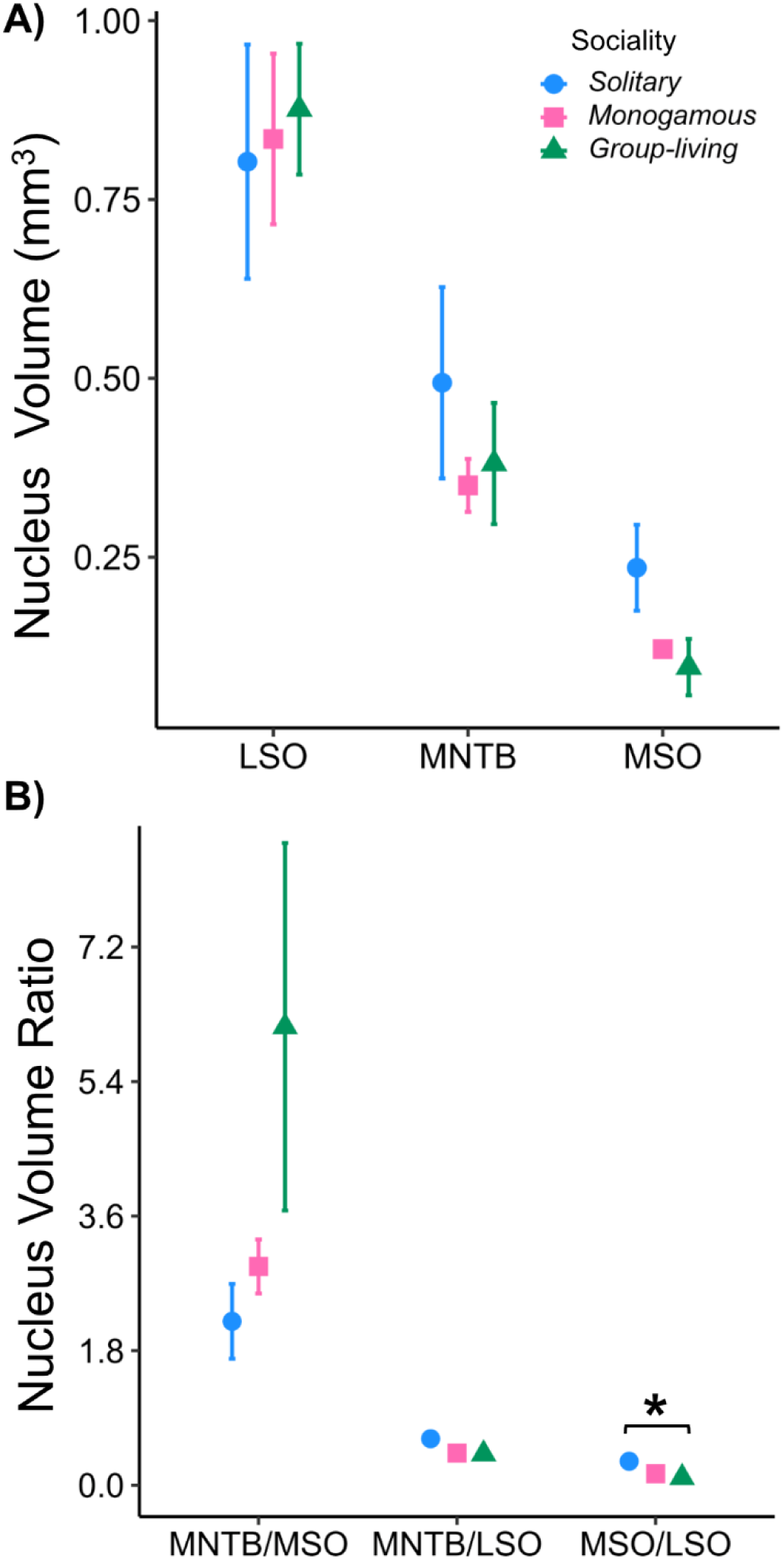
Auditory brainstem nucleus and relative nucleus volume ratios across social groups. A) Mean volumes of the LSO, MNTB, MSO, across solitary, monogamous, and group-living species. B) Relative volumes of the MNTB, MSO, and LSO expressed as ratios to LSO and MSO volume (MNTB/LSO, MNTB/MSO, and MSO/LSO) across social groups (blue circle = solitary, pink squared = monogamous, and green triangles (group-living). Values display represent means and standard errors for each social group. Significant species differences were observed in both MNTB and MSO volume across species, but not LSO volume. No significant difference was detected in MNTB, MSO, and LSO volume among social groups. Significant differences in MSO/LSO ratio between solitary and group-living species. Asterisks denoted statistically significant with * p < 0.05.

## DISCUSSION

Understanding the neural mechanisms that support the diversity of animal social behavior requires examining not only higher-order brain region neuromodulatory circuits but also early sensory circuits that facilitate intraspecific communication and social interaction. In the present work, we compared the number of OT and AVP immunoreactive puncta within three principal nuclei of the auditory brainstem (LSO, MSO, MNTB) across six wild-caught rodent species exhibiting different forms of social behavior strategies. We also examined associations between sociality and total brain mass, total brain volume, and the volume of brainstem nuclei volume among the studied rodent species. We found significant species and sociality groups differences in OT and AVP immunoreactive puncta, with group-living species generally exhibiting higher OT and AVP puncta counts than monogamous and solitary species in the LSO and MNTB. In contrast, OT puncta count in the MSO did not differ among species or sociality groups, whereas group-living species exhibited higher AVP puncta counts in the MSO than monogamous and solitary species. However, the MSO was relatively small in *M. musculus*, and the sampled field of view may have included adjacent brain regions for this species. Therefore, the MSO findings should be interpreted with caution. Brain mass differed significantly among both species and social groups, while total brain volume differed significantly among species but was not associated with sociality. Notably, the group-living *M. musculus* exhibited a relatively small total brain volume compared with several of the other species examined, indicating that differences in auditory neuropeptide expression were not simply attributable to overall brain size. These findings suggest that variation in OT and AVP immunoreactive puncta within early auditory circuits is associated with differences in social organization across rodent species and may reflect differences in the neurochemical modulation of auditory processing involved in socially relevant communication.

This study revealed significant overall species effects on OT and AVP immunoreactive puncta counts in the LSO and MNTB of the auditory brainstem. However, these overall effects were not supported by significant differences between individual species in Tukey-adjusted pairwise comparisons. Previous studies have reported species differences in OT and AVP neurons, receptors, fibers, and puncta in the forebrain and midbrain regions across mammals (Hashimoto et al., 1985; Rogers Flattery et al., 2022; Xu et al., 2010). For instance, Xu and colleagues (2010) found that greater long-tailed hamsters (*Tscherskia triton*) exhibited lower density of OT immunoreactive cells in the medial amygdala and the medial preotic area, as well as had lower density of AVP immunoreactive cells in the lateral hypothalamus, compared to Brandt’s voles (*Lasiopodomys brandtii*). In addition, OT receptors are detected in reward-related regions such as the ventral pallidum and nucleus accumbens in humans (*Homo sapiens*) but are not present in these areas in chimpanzees (*Pan troglodytes*), suggesting notable interspecific variation in the distribution of these neuropeptide hormone receptors (Rogers Flattery et al., 2022). These findings support substantial interspecific variation in the distribution of OT and AVP throughout the mammalian brain. Such variation may reflect differences in the organization and functional demands of neural circuits involved in social behavior, as well as species-specific ecological and evolutionary histories. Thus, rather than exhibiting a uniform pattern across taxa, OT and AVP systems may be differentially organized across brain regions on ways that correspond to the diverse social and ecological strategies of mammals.

As expected, group-living species exhibited a higher number of OT immunoreactive puncta than both monogamous and solitary-living species across the LSO and MNTB nuclei. However, no significant differences in OT puncta were detected among groups in the MSO. This partially supports our hypothesis that social species would exhibit greater OT puncta expression within auditory brainstem nuclei compared to other social group types. However, previous studies examining other brain regions have not consistently found differences in OT distribution across species with varying social organizations. For instance, OT immunoreactive cells in the lateral hypothalamus and medial preotic area were similar among monogamous and polygamous vole species (Shapiro & Insel, 1992; Wang et al., 1996). Similarly, OT receptor densities in the medial preotic area were comparable between the social *Ctenomys sociabilis* and the solitary *Ctenomys haigi* (Beery et al., 2008). Accordingly, differences in OT immunoreactive puncta among social groups in this present investigation may reflect variation in broader social lifestyle strategies rather than species-related specific behaviors. While the functional significance of these neuropeptide differences within the LSO, MSO, and MNTB nuclei remain unclear, these nuclei are well established as key components of early auditory processing pathways. Therefore, variation in oxytocinergic puncta signaling within these nuclei may indicate that neuromodulatory systems associated with social behavior interact with auditory circuits at early stages of sensory processing. This raises the possibility that such modulation could influence the processing of socially relevant acoustic signals, although this hypothesis needs to be investigated.

Our previous work has demonstrated that group-living rodents exhibit greater sensitivity in hearing thresholds and enhanced binaural hearing compared to monogamous and solitary species (McCullagh et al., 2025). Because the LSO, MSO, and MNTB are key nuclei involved in binaural integration and sound localization (Haragopal & Winters, 2023; Joseph et al., 2026; Joseph et al., 2025; Middlebrooks, 2015; Tollin, 2003), the higher number of AVP and OT-puncta observed in group-living species may suggest a potential neuromodulatory role for AVP and OT within early auditory pathways. More broadly, a growing body of work across vertebrates demonstrates that steroid, OT, AVP, and vasotocin hormones modulate both the production of vocal signals and the processing of socially relevant auditory cues (Penna et al., 1992; Boyd 1994; Klomberg et al., 2000; Caras et al. 2010). For example, in amphibians and birds, steroid and arginine vasotocin (the homolog of vasopressin) hormones regulate vocal communication and behavioral responses to conspecific calls (Maler et al., 1995; Propper and Dixon 1997; Semsar et al., 1998; Caras et al., 2010; Kim et al., 2010; Caras & Remage-Healey 2016). Similarly, in rodents, OT and AVP signaling within forebrain and midbrain regions have been linked to influence vocal signal production (Floody et al., 1998; Charlton et al., 2019). In this context, the present study extends this framework by demonstrating variation in oxytocinergic and vasopressinergic immunoreactive puncta within early auditory brainstem nuclei across rodent species with differing social organization strategies. While prior work has been largely devoted to vocal production or higher brain-region processing, our work suggests that social neuromodulatory systems may also influence auditory processing at the level of early sensory circuits. To our knowledge, this is the first study to quantify OT and AVP-immunoreactive puncta within the LSO, MSO, and MNTB of rodents differing in social behavior. Future investigations should therefore characterize the distribution of OT and AVP receptors within these brainstem nuclei and test whether variation in OT and AVP receptors or puncta numbers influence neural activity and auditory computation within early sensory circuits.

Likewise, social group differences were also detected in AVP immunoreactive puncta across the studied nuclei. Group-living rodents exhibited higher numbers of AVP immunoreactive puncta in the LSO, MNTB, and MSO regions of the auditory brainstem compared to solitary rodents. However, there was no difference in the numbers of AVP puncta across measured auditory brainstem regions between monogamous and solitary species. While AVP immunoreactive puncta in these nuclei has not been the focus of prior investigation, group differences in AVP neuron densities have been documented in the medial preotic area, the lateral hypothalamus, and the bed nucleus of the stria terminalis of mice and rats (Wang, 1995). Similarly, the relative density of AVP immunoreactive cells differ in other brain regions among taxa that exhibit different social living styles (Wang, 1995; Xu et al., 2010). For instance, social Brandt’s voles had higher densities of AVP immunoreactive cells in the lateral hypothalamus than solitary greater long-tailed hamsters (Xu et al., 2010). Similar patterns have been observed between monogamous prairie voles (*Microtus ochrogaster*) and promiscuous meadow voles (*Microtus pennsylvanicus*), in which prairie voles displayed higher expression of AVP cells in the bed nucleus of the stria terminalis than meadow voles (Wang, 1995). However, in other brain regions, opposite patterns have been detected among species differing in sociality. Solitary white-footed mice (*Peromyscus leucopus*) had more AVP immunoreactive cells in the bed nucleus of the stria terminalis compared to monogamous California mice (*Peromyscus californicus*), while Greater long-tailed hamsters had higher AVP immunoreactive cells in the medial preotic area than social Brandt’s voles (Bester-Meredith et al., 1999; Xu et al., 2010). The reasons for these inconsistent results remain unclear. However, AVP involves diverse physiological and behavioral functions in the brain, not all of which are directly related to social behavior, and its neural effects can vary across brain regions and species (Cambell 2010; Xue et al., 2010).

Conventionally, brain size has been used as a proxy for cognitive capacity in tests of the social brain hypothesis, which proposes that increased social complexity is associated with larger brains (Dunbar, 2009). Evidence for this hypothesis has largely come from interspecific comparisons, although findings have been mixed across taxa. Positive associations between social behavior and brain size have been reported in primates (Dunbar, 1998), ungulates (Shultz & Dunbar, 2006), fishes (Ma et al., 2025), carnivores, and insectivores (Dunbar & Bever, 1998; Pérez-Barbería et al., 2007). In contrast, comparative studies in birds (Hardie & Cooney, 2023) and rodents (Kverková et al., 2018) have failed to identify consistent associations between sociality and brain size and, in some cases, have found larger brains in less social species. These contrasting findings highlight the complexity of brain evolution and the potential influence of ecological and life-history factors that covary with social organization, including foraging strategy, habitat, and developmental mode (Hardie & Cooney, 2023; Kverková et al., 2018; Xu et al., 2010). For instance, Hardie and Cooney (2023) found that developmental mode and foraging strategy were more relevant for predicting brain size in birds than social organization, whereas Kverková and Colleagues (2018) reported larger brain in solitary African mole-rats than in eusocial species. Our findings similarly provide limited support for a simple association between sociality and overall brain size. Although total brain volume differed among species, it did not differ among social groups. Brain mass, however, differed significantly among both species and social groups, with solitary species exhibiting heavier brain mass than monogamous species, but not differing significantly from group-living species. Moreover, body mass was significantly associated with brain mass and brain volume when species were compared across social groups, emphasizing the importance of accounting for body size when interpreting interspecific variation in brain morphology. Together, these results indicate that sociality alone does not consistently explain variation in global brain morphology among rodent species examined here. Instead, differences in social behavior may be more closely associated with variation in the organization and modulation of species-specific neural circuits, including oxytocinergic and vasopressinergic systems, that support social communication and sensory processing.

Our brainstem nuclei volumetric analyses revealed significant interspecific differences in MNTB and MSO volume, whereas LSO volume remained relatively conserved across species, suggesting that variation in size of individual auditory nuclei is driven primarily by species-specific characteristics rather than sociality of the studied species. Consistent with this interpretation, no differences in the absolute volumes of the LSO, MSO, or MNTB were detected when species were grouped by social organization. To account for interspecific differences in nucleus sizes, we also compared the MNTB/LSO, MNTB/MSO and MSO/LSO volume ratios. Similar to previous studies (E. A. McCullagh et al., 2022), ratio-based analyses allowed us to assess the relative organization of auditory brainstem nuclei independent of overall size. Although the MNTB/LSO and MNTB/MSO ratio did not differ across species or social groups, the MSO/LSO ratio was smaller in group-living than solitary species, indicating a proportionally smaller MSO relative to the LSO in group-living rodents. Together, these findings suggest that sociality may be associated with differences in the relative organization of auditory brainstem nuclei rather than their absolute volumes.

There are numerous limitations that should be considered when interpreting these results. First, the use of wild-caught rodents introduces inevitable biological variability in age, developmental history, reproductive status, and prior environmental exposure or life experiences, all of which may play a role in differences in OT and AVP immunoreactive puncta number reported here. Second, the immunohistochemical technique used in this work solely quantified the number of puncta, but does not explicitly provide information about neuromodulatory cells, receptor activation, and synaptic function within these brainstem nuclei. Therefore, the present variable (puncta counts) should be interpreted as anatomical correlations of neuromodulator signaling rather than direct measures of functional neuronal activity within the LSO, MSO, and MNTB. Third, while statistical comparisons were performed across species, phylogenetic relatedness may partially influence the observed pattern shown in this study, as closely related taxa tend to have conserved neuroanatomical traits independent of social organization. However, phylogeny was not accounted for in any data analyses because the sample size (N = 6) was too small to support robust phylogenetically informed analyses. Finally, although the correlation between social status and neuromodulator immunoreactive puncta numbers is evident across the studied brainstem nuclei, the pathway of these two neuropeptides hormones from the hypothalamus to the auditory brainstem, as well as the developmental mechanism underlying these relationships remain unclear. Future investigations should extend these findings by integrating comparative neuroanatomy with physiological and behavioral experiments. Combining OT and AVP mapping with physiological measures such as auditory brainstem responses, *ex vivo* and *in vitro* electrophysiology techniques would aid in determining whether variation in the density of OT and AVP neurons influences auditory sensitivity, temporal precision, or binaural processing within the auditory brainstem. In addition, behavioral experiments using socially relevant acoustic stimuli, such as pup calls or conspecific vocalization playbacks, could further test whether species within each social group that exhibit higher peptide densities show enhanced neural or behavioral responsiveness to signals. Future studies should also use more spatially restricted sampling methods, particularly in species with a small or poorly delineated MSO, to minimize the potential contribution of adjacent brain regions to measured immunoreactivity. Expanding the analysis to include both sexes would also help determine whether the observed patterns of OT and AVP immunoreactivity are consistent across males and females. Finally, incorporating phylogenetic comparative analyses and expanding sampling across additional rodent clades, and other vertebrates, will be essential for clarifying the evolutionary contributions of shared ancestry, ecological pressures, and social complexity to the organization of early auditory circuits.

## CONCLUSION

The results of this study provide evidence that social lifestyle strategies are a strong predictor of OT and AVP immunoreactive puncta within nuclei of the auditory brainstem. Despite no differences in total brain volume across social categories, OT and AVP puncta density within the LSO, MSO, and MNTB differ considerably among social groups, with social species consistently exhibiting higher puncta counts across measured nuclei. These results suggest that early circuits of auditory processing may be important targets of social selection in rodents, and that neuromodulatory signals may shape the neural representation of socially relevant acoustic information. By incorporating comparative neuroanatomy, this study advances our understanding of the distribution of OT and AVP puncta within the LSO, MNTB, and LSO nuclei of the auditory brainstem across rodents that exhibit different social behavior organizations.

## ACKNOWLEGEMENTS

We thank the personnels of Packsaddle, James Collins, and Sandy Sanders wildlife management areas, Sheena Parsons, Benny Farrar, Marie Stone, and Marcus Thibodeau for housing and permission to collect rodents in our sampling locations. We also thank Sarah Hobbs and Naleyshka Colon-Rivera for helping in slicing brain and immunohistochemistry staining to complete this work. This work was partially supported by an NSF CAREER award to Elizabeth A. McCullagh (244070).

