## supplemental figures and tables for "Oxytocin and Vasopressin Immunoreactivity Differs Across Auditory Brainstem Nuclei in Rodents with Distinct Social Systems"

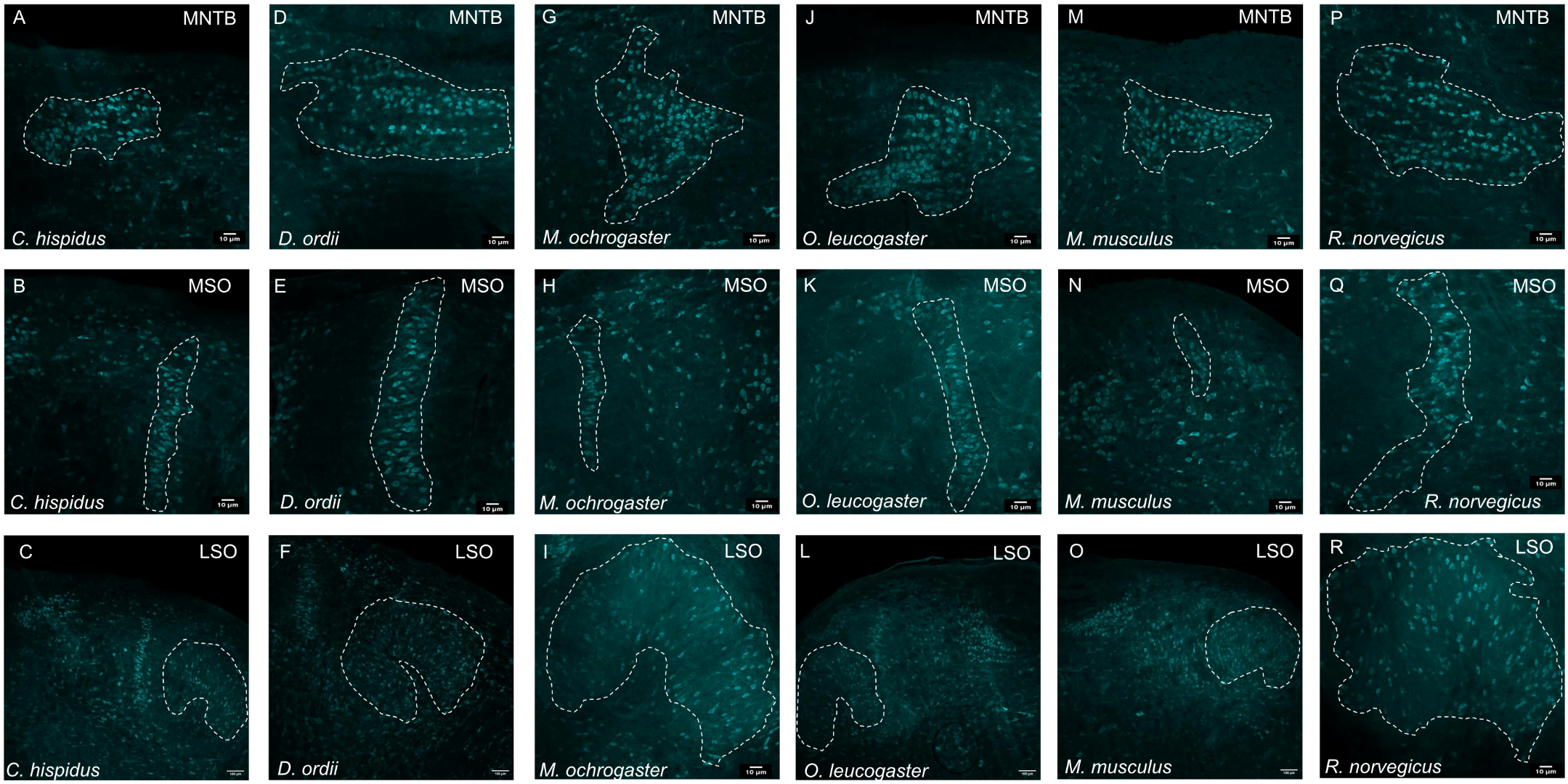

**Supplementary Figure 1:** Representative Nissl staining of MNTB, MSO, and LSO of each studied species. Each image represents a 10x image (scale at 100 microns) of the MNTB (**A, D, G, J, M, P**), MSO (**B, E, H, K, N, Q**) and LSO (**C, F, I, L, O, R**). White outlines indicate the boundaries of each nucleus used for volume measurements.

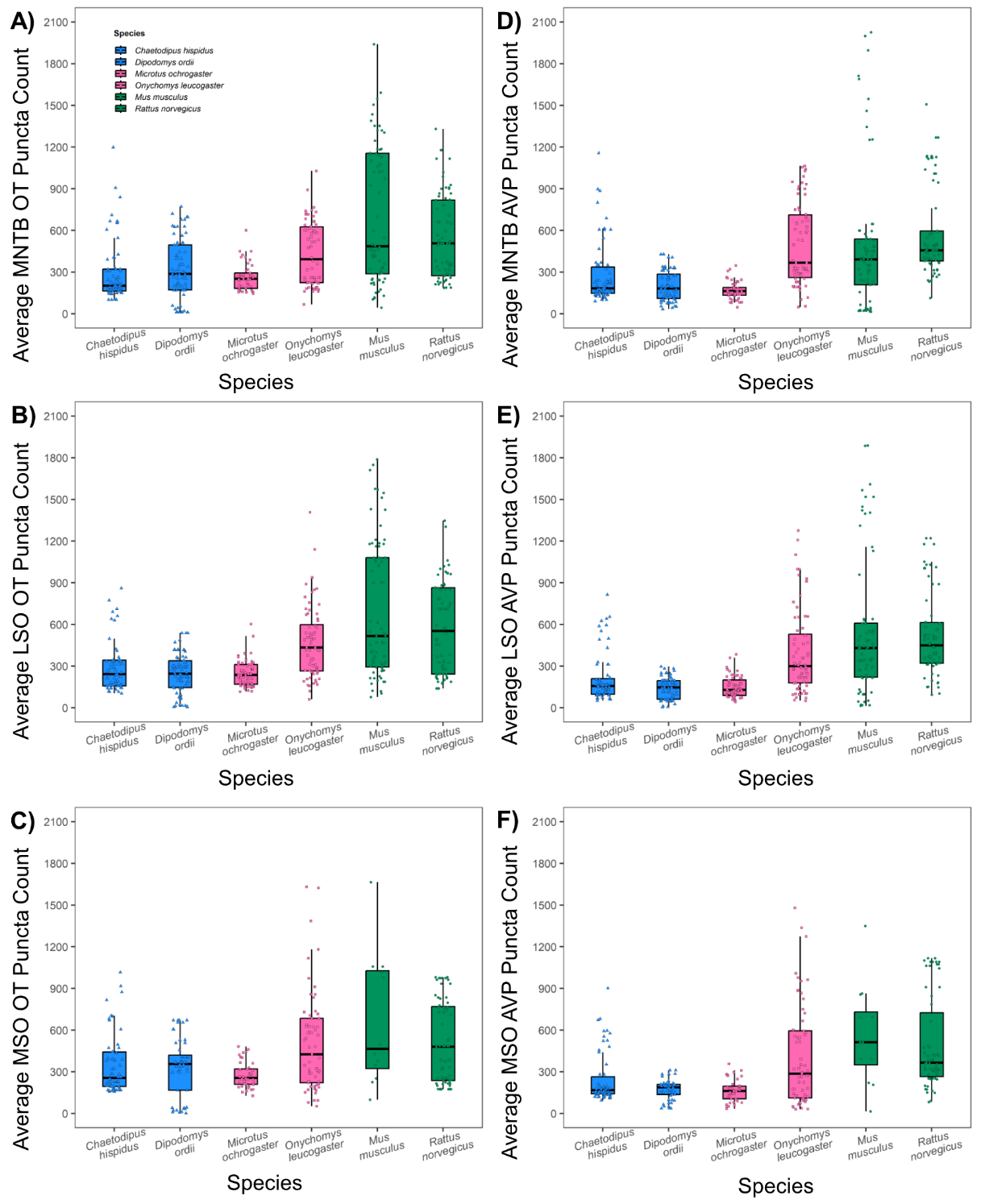

**Supplementary Figure 2:** Oxytocin (OT) and vasopressin (AVP) immunoreactive puncta counts across species in the MNTB, LSO, and MSO. A-C, Average OT immunoreactive puncta counts across species (solitary species in dodger blue, monogamous species in hot pink, and group-living species in green) in the MNTB (A), LSO (B) and MSO (C), respectively. D-F, Average AVP-immunoreactive puncta counts across species in the MNTB (D), LSO (E), and MSO (F), respectively. Boxplots show the median and interquartile range; whiskers extend to 1.5 times the interquartile range. Individual point represent puncta count measurements. Significant differences in OT and AVP puncta counts were detected among species.

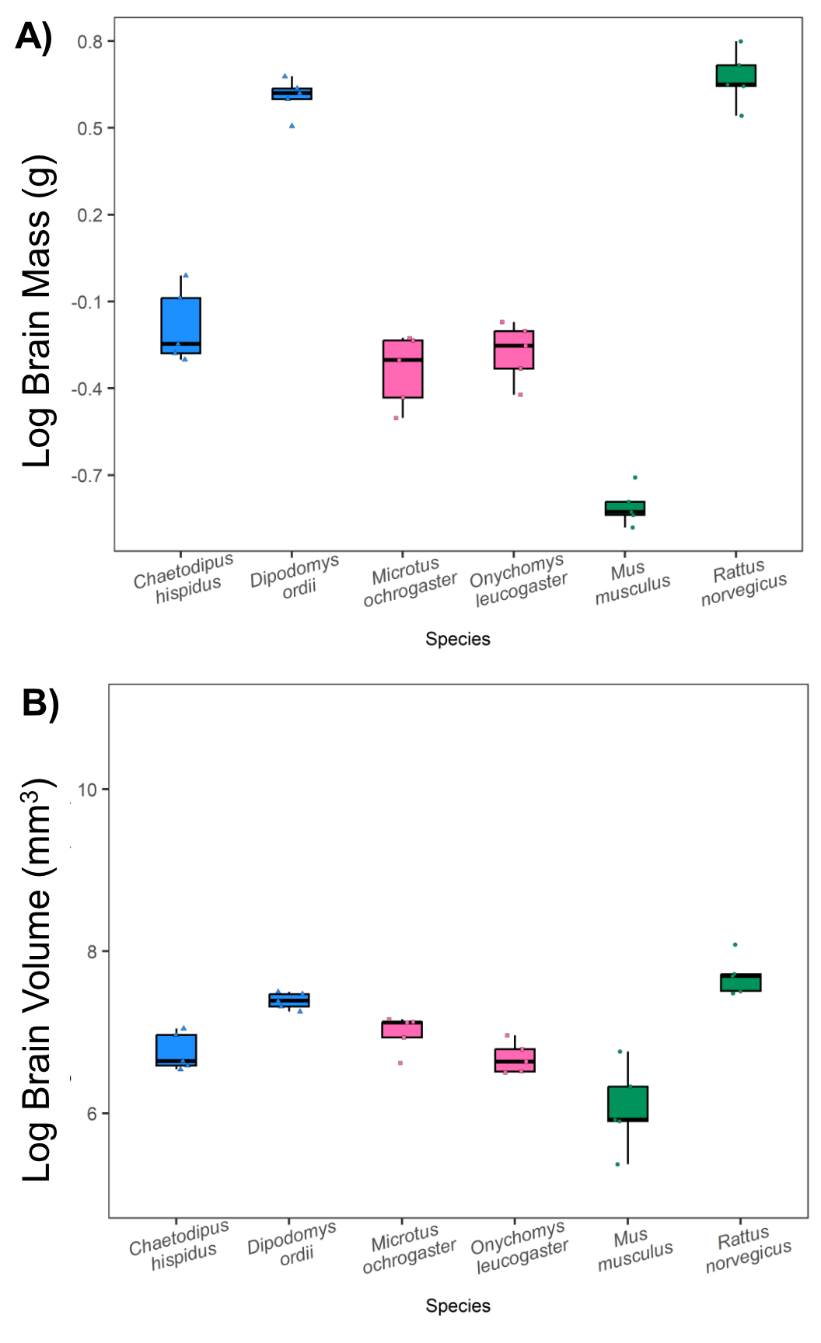

**Supplementary Figure 3:** Log-transformed brain mass and brain volume across species. A) Log-transformed brain mass across species B) Log-transformed brain volume across species. Each point represents an individual animal and is colored and shaped according to its specie-specific sociality classification: solitary species are represented by blue triangles, monogamous species by pink squares and group-living species by green circles. Boxplots show the median and interquartile range; whiskers extend to 1.5 times the interquartile range. Individual point represent log-transformed brain mass and volume for each species. Significant differences in log-transformed brain mass and brain volume were detected across species.

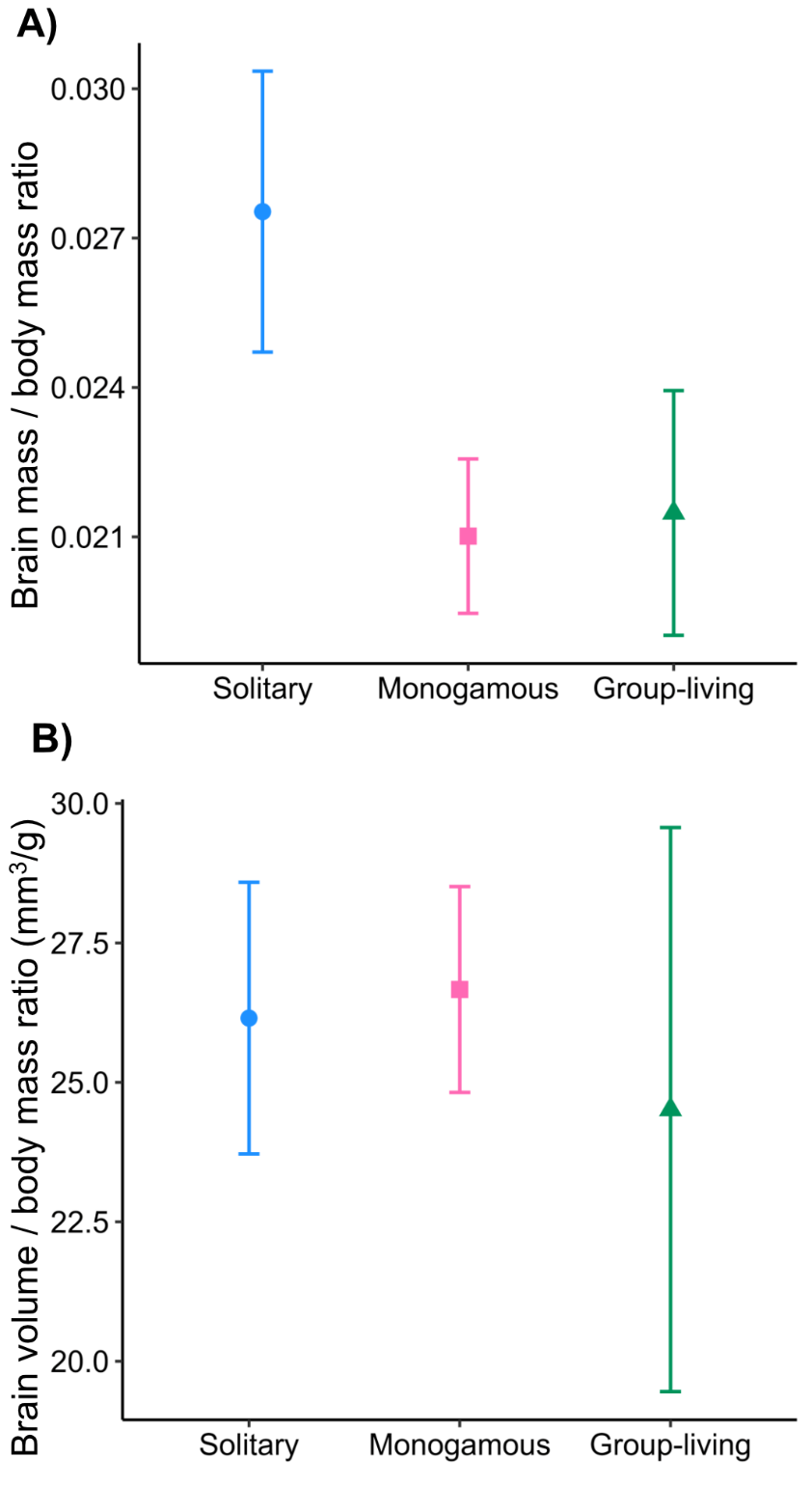

**Supplementary Figure 4:** Brain volume / brain body mass ratio and brain mass / body mass ratio across social groups. A) Mean brain mass / body mass ratios across social groups B) Mean brain volume / brain mass across social groups. No significant sociality differences were observed in both the brain mass / body mass ratio and brain volume / body mass ratio..

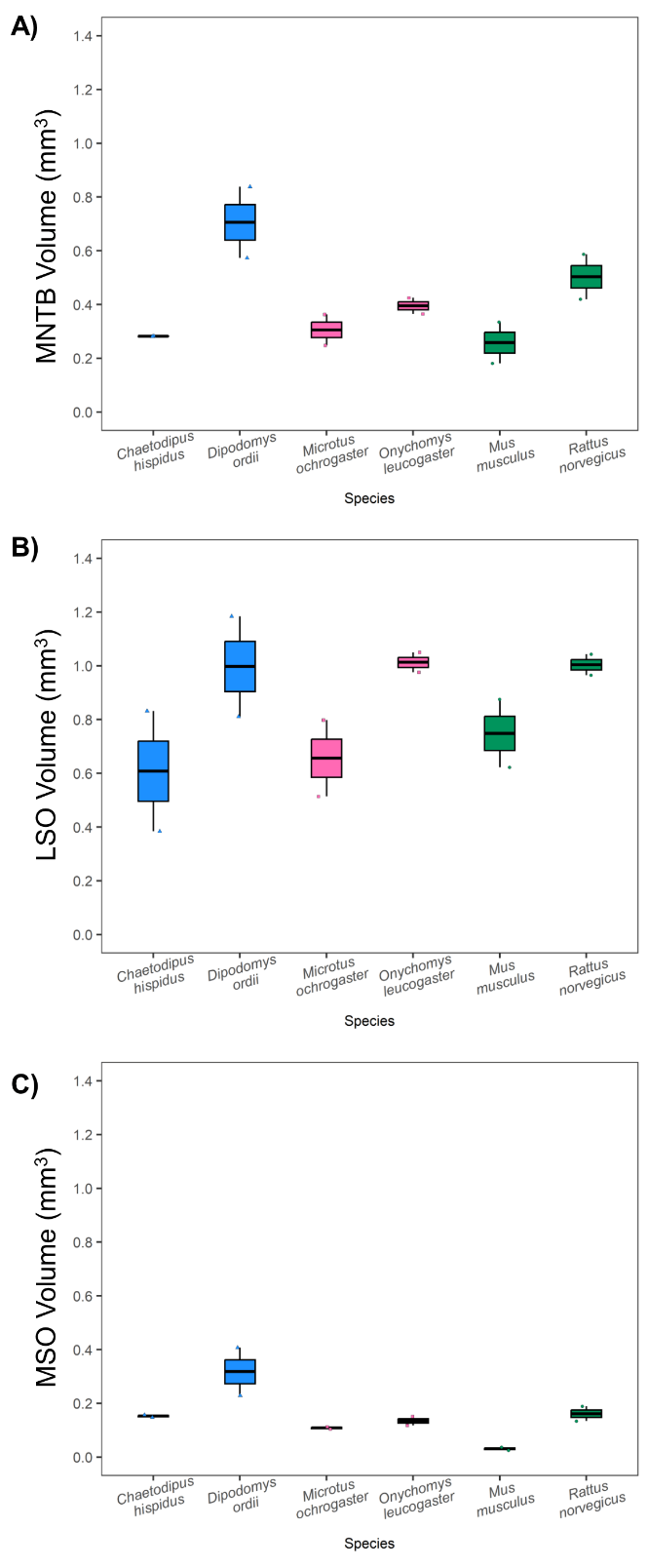

**Supplementary Figure 5:** Auditory brainstem nucleus volume across species. A) Mean volumes of the MNTB across species, B) Mean volumes of the LSO across species, C) Mean volumes of the MSO across species. Boxplots show the median and interquartile range; whiskers extend to 1.5 times the interquartile range. Individual point represent nuclei volume. Significant species differences were observed in both MNTB and MSO volume across species.

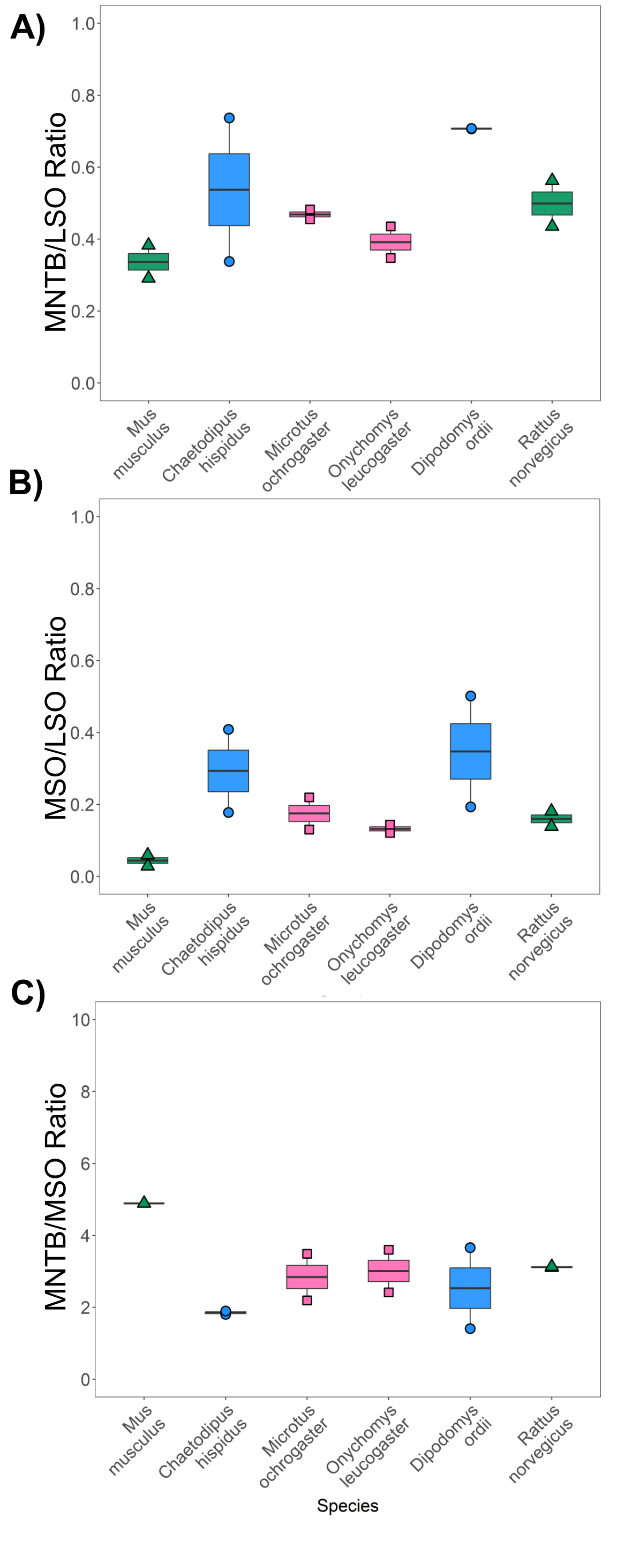

**Supplementary Figure 6:** Auditory brainstem nucleus volume ratio across species. A) Mean MNTB/ LSO volume ratios across species, B) Mean MSO/LSO volume ratios across species. C) Mean MNTB / MSO volume ratios across species. Boxplots show the median and interquartile range; whiskers extend to 1.5 times the interquartile range. Individual point represent the ratio value. No significant species differences were observed in both the MNTB/LSO and MSO/LSO volume ratios across species.

**Table 1:** Pairwise comparisons of log-transformed brain mass across species. Value displayed represent the estimate, standard error (SE), domain of definition (Df), t-ratio value, and the p-value.

| **Log brain volume Pairwise Contrasts Across Species** | **Estimate** | **SE** | **Df** | **t-ratio** | **p-value** |
| --- | --- | --- | --- | --- | --- |
| *Chaetodipus hispidus – Dipodomys ordii* | - 0.732 | 0.06 | 23 | -10.610 | **< 0.0001** |
| *Chaetodipus hispidus - Microtus ochrogaster* | 0.160 | 0.06 | 23 | 2.659 | 0.1222 |
| *Chaetodipus hispidus - Mus musculus* | 0.547 | 0.07 | 23 | 7.393 | **< 0.0001** |
| *Chaetodipus hispidus - Onychomys leucogaster* | 0.085 | 0.06 | 23 | 1.410 | 0.7210 |
| *Chaetodipus hispidus - Rattus norvegicus* | -0.728 | 0.09 | 23 | -7.539 | **< 0.0001** |
| *Dipodomys ordii - Microtus ochrogaster* | 0..892 | 0.06 | 23 | 13.220 | **< 0.0001** |
| *Dipodomys ordii - Mus musculus* | 1.279 | 0.09 | 23 | 13.109 | **< 0.0001** |
| *Dipodomys ordii - Onychomys leucogaster* | 0.817 | 0.07 | 23 | 11.549 | **< 0.0001** |
| *Dipodomys ordii - Rattus norvegicus* | 0.010 | 0.07 | 23 | 0.143 | 1.000 |
| *Microtus ochrogaster - Mus musculus* | 0.386 | 0.07 | 23 | 5.096 | **0.0005** |
| *Microtus ochrogaster - Onychomys leucogaster* | -0.075 | 0.06 | 23 | -1.243 | 0.811 |
| *Microtus ochrogaster - Rattus norvegicus* | -0.882 | 0.09 | 23 | -9.455 | **< 0.0001** |
| *Mus musculus - Onychomys leucogaster* | -0.462 | 0.07 | 23 | -6.413 | **< 0.0001** |
| *Mus musculus - Rattus norvegicus* | -1.268 | 0.13 | 23 | -9.611 | **< 0.0001** |
| *Onychomys leucogaster - Rattus norvegicus* | -0.806 | 0.09 | 23 | -8.196 | **< 0.0001** |

**Table 2:** Pairwise comparisons of log-transformed brain volume across species. Value displayed represent the estimate, standard error (SE), domain of definition (Df), t-ratio value, and the p-value.

| **Log brain volume Pairwise Contrasts Across Species** | **Estimate** | **SE** | **Df** | **t-ratio** | **p-value** |
| --- | --- | --- | --- | --- | --- |
| *Chaetodipus hispidus – Dipodomys ordii* | - 0.577 | 0.21 | 23 | -2.766 | 1.000 |
| *Chaetodipus hispidus - Microtus ochrogaster* | -0.231 | 0.18 | 23 | -1.269 | 0.798 |
| *Chaetodipus hispidus - Mus musculus* | 0.633 | 0.22 | 23 | 2.829 | 0.088 |
| *Chaetodipus hispidus - Onychomys leucogaster* | 0.069 | 0.18 | 23 | 0.381 | 0.998 |
| *Chaetodipus hispidus - Rattus norvegicus* | -0.856 | 0.29 | 23 | -2.852 | 0.084 |
| *Dipodomys ordii - Microtus ochrogaster* | 0.346 | 0.20 | 23 | 1.694 | 0.548 |
| *Dipodomys ordii - Mus musculus* | 1.211 | 0.29 | 23 | 4.101 | **0.005** |
| *Dipodomys ordii - Onychomys leucogaster* | 0.647 | 0.21 | 23 | 3.022 | 0.059 |
| *Dipodomys ordii - Rattus norvegicus* | -0.248 | 0.22 | 23 | -1.131 | 0.863 |
| *Microtus ochrogaster - Mus musculus* | 0.865 | 0.23 | 23 | 3.766 | **0.011** |
| *Microtus ochrogaster - Onychomys leucogaster* | 0.301 | 0.18 | 23 | 1.643 | 0.580 |
| *Microtus ochrogaster - Rattus norvegicus* | -0.594 | 0.28 | 23 | -2.107 | 0.318 |
| *Mus musculus - Onychomys leucogaster* | -0.564 | 0.21 | 23 | -2.587 | 0.141 |
| *Mus musculus - Rattus norvegicus* | -1.460 | 0.40 | 23 | -3.654 | **0.014** |
| *Onychomys leucogaster - Rattus norvegicus* | -0.896 | 0.29 | 23 | -3.007 | 0.061 |

**Table 3:** Pairwise comparisons of MNTB volume across species. Value displayed represent the estimate, standard error (SE), domain of definition (Df), t-ratio value, and the p-value.

| **MNTB Volume Pairwise Contrasts Across Species** | **Estimate** | **SE** | **Df** | **t-ratio** | **p-value** |
| --- | --- | --- | --- | --- | --- |
| *Chaetodipus hispidus – Dipodomys ordii* | -0.423 | 0.107 | 6 | -3.940 | 0.052 |
| *Chaetodipus hispidus - Microtus ochrogaster* | -0.023 | 0.107 | 6 | -1.219 | 0.999 |
| *Chaetodipus hispidus - Mus musculus* | 0.024 | 0.107 | 6 | 0.223 | 0.999 |
| *Chaetodipus hispidus - Onychomys leucogaster* | -0.113 | 0.107 | 6 | -1.051 | 0.884 |
| *Chaetodipus hispidus - Rattus norvegicus* | -0.221 | 0.107 | 6 | -2.061 | 0.408 |
| *Dipodomys ordii - Microtus ochrogaster* | 0.400 | 0.107 | 6 | 3.721 | 0.066 |
| *Dipodomys ordii - Mus musculus* | 0.447 | 0.107 | 6 | 4.163 | **0.041** |
| *Dipodomys ordii - Onychomys leucogaster* | 0.310 | 0.107 | 6 | 2.889 | 0.166 |
| *Dipodomys ordii - Rattus norvegicus* | 0.202 | 0.107 | 6 | 1.879 | 0.488 |
| *Microtus ochrogaster - Mus musculus* | 0.047 | 0.107 | 6 | 0.442 | 0.996 |
| *Microtus ochrogaster - Onychomys leucogaster* | -0.089 | 0.107 | 6 | -0.833 | 0.950 |
| *Microtus ochrogaster - Rattus norvegicus* | -0.198 | 0.107 | 6 | -1.842 | 0.506 |
| *Mus musculus - Onychomys leucogaster* | -1.137 | 0.107 | 6 | -1.274 | 0.789 |
| *Mus musculus - Rattus norvegicus* | -0.245 | 0.107 | 6 | -2.284 | 0.324 |
| *Onychomys leucogaster - Rattus norvegicus* | -0.108 | 0.107 | 6 | -1.009 | 0.899 |

**Table 4:** Pairwise comparisons of MSO volume across species. Value displayed represent the estimate, standard error (SE), domain of definition (Df), t-ratio value, and the p-value.

| **MSO Volume Pairwise Contrasts Across Species** | **Estimate** | **SE** | **Df** | **t-ratio** | **p-value** |
| --- | --- | --- | --- | --- | --- |
| *Chaetodipus hispidus – Dipodomys ordii* | -0.165 | 0.055 | 6 | -3.017 | 0.144 |
| *Chaetodipus hispidus - Microtus ochrogaster* | 0.044 | 0.055 | 6 | 0.802 | 0.957 |
| *Chaetodipus hispidus - Mus musculus* | 0.121 | 0.055 | 6 | 2.215 | 0.348 |
| *Chaetodipus hispidus - Onychomys leucogaster* | 0.018 | 0.055 | 6 | 0.328 | 0.999 |
| *Chaetodipus hispidus - Rattus norvegicus* | -0.009 | 0.055 | 6 | -0.164 | 1.000 |
| *Dipodomys ordii - Microtus ochrogaster* | 0.209 | 0.055 | 6 | 3.819 | 0.059 |
| *Dipodomys ordii - Mus musculus* | 0.287 | 0.055 | 6 | 5.232 | **0.014** |
| *Dipodomys ordii - Onychomys leucogaster* | 0.183 | 0.055 | 6 | 3.345 | 0.099 |
| *Dipodomys ordii - Rattus norvegicus* | 0.156 | 0.055 | 6 | 2.853 | 0.173 |
| *Microtus ochrogaster - Mus musculus* | 0.077 | 0.055 | 6 | 1.413 | 0.722 |
| *Microtus ochrogaster - Onychomys leucogaster* | -0.026 | 0.055 | 6 | -0.474 | 0.995 |
| *Microtus ochrogaster - Rattus norvegicus* | -0.053 | 0.055 | 6 | -0.966 | 0.913 |
| *Mus musculus - Onychomys leucogaster* | -1.103 | 0.055 | 6 | -1.887 | 0.485 |
| *Mus musculus - Rattus norvegicus* | -0.131 | 0.055 | 6 | -2.379 | 0.292 |
| *Onychomys leucogaster - Rattus norvegicus* | -0.0.27 | 0.055 | 6 | -0.492 | 0.995 |
